# *OsNLP4* is required for nitrate assimilation gene expressions and nitrate-dependent growth in rice

**DOI:** 10.1101/2020.03.16.993733

**Authors:** Mengyao Wang, Takahiro Hasegawa, Makoto Hayashi, Yoshihiro Ohmori, Kenji Yano, Shota Teramoto, Takehiro Kamiya, Toru Fujiwara

**Affiliations:** The Laboratory of Plant Nutrition and Fertilizers, Graduate School of Agricultural and Life Sciences, The University of Tokyo, Tokyo, 1138657, Japan; RIKEN Center for Sustainable Resource Science, Kanagawa, 2300045, Japan

**Keywords:** nitrate, rice NLP family, growth, nitrate-related genes, nitrate assimilation

## Abstract

In plants, nitrate is important nutrient and signaling molecule that modulates the expression of many genes and regulates growth. In paddy grown rice, nitrogen is mostly supplied in the form of ammonium, but nitrate also shares substantial portion of available nitrogen. A number of nitrogen transporters and nitrate assimilation enzymes have been identified and functionally characterized. However, little is known about the nitrate sensor system and regulatory mechanisms of these nitrate related genes. In recent years, NIN-like proteins (NLPs) have been described as key transcription factors of nitrogen responses in *Arabidopsis thaliana*. But the functions of OsNLPs in rice are still elusive. Here we report the characterization of *OsNLP4* to reveal its function in rice. Growths of *OsNLP4* knockout mutants were reduced under the nitrate supply, but not under ammonium supply. The mRNA accumulation of genes involved in nitrate assimilation were declined significantly and nitrate uptake rate and nitrate reductase activity were also impaired in the mutants. Using rice protoplast transient expression system, OsNLP4-GFP fusion was localized to nucleus irrespective of nitrate conditions. *OsNLP4* was also required for normal yield under paddy field conditions. We propose the *OsNLP4* is essential for regulation of genes involved in nitrate assimilation and nitrate-dependent growth in rice.

**One sentence summary:** The *osnlp4* mutants exhibit abnormal nitrate response and poor growth under nitrate supply and in paddy field conditions.

## Introduction

Nitrogen (N) is the most important macronutrient for plant growth and development. Nitrate (NO_3_^−^) and ammonium (NH_4_^+^) are the major sources of inorganic N taken up by the roots of higher plants. Nitrate is generally present at higher concentrations than those of ammonium in the soil solution under aerobic conditions. In flooded paddy soils, ammonium is the abundant form of nitrogen than nitrate. In both cases N levels in soil fluctuates by rainfalls and addition of N sources etc. and it is necessary for plants to adapt to changing N conditions. Land plants display phenotypic plasticity and reprogram gene expression in response to nitrogen availability (Zhang and Forde 2000; Stitt 1999; Forde 2002; Sasakawa and Yamamoto 1978). In general, elongation of seminal and lateral roots of rice seedlings is inhibited by high concentrations of ammonium, but not by high nitrate levels (Hirano et al. 2008). For nitrate, it is also known to affect expression of a wide spectrum of genes, including not only genes related to nitrate transport and assimilation, but also genes involved in other metabolic pathways and plant development (Wang et al. 2003; Wang et al. 2007; Crawford 1995). For instance, *AtNIA1* and *AtNIA2* (nitrate reductase genes) are induced by nitrate and determine the levels of NO production and ABA-induced stomatal closure in *A. thaliana* (Desikan et al. 2002; Chen et al. 2016; Zhao et al. 2016). In *A. thaliana*, these responses are mediated by several factors including the nitrate transceptor NRT1.1 (Ho et al. 2009), two kinases CIPK8/23 (Ho et al. 2009; Hu et al. 2009) and the transcription factors NLP6/7/8 (Yanagisawa 2014; Yan et al. 2016; Castaings et al. 2009), LBD37-LBD39 (Rubin et al. 2009), TCP20 (Guan et al. 2017; Guan et al. 2014) and SPL9 (Krouk et al. 2010b). In rice, the functional characterization of a number of nitrate transporter genes have been reported. Such as *OsNRT2.3a*, one of the nitrate transporter genes in rice induced under nitrate supply, plays a key role in long-distance nitrate transport from root to shoot at low nitrate supply level (Tang et al. 2012). OsNPF7.3, a member of the nitrate transporter 1/peptide transporter family, enhances nitrogen allocation and increase grain yield in rice (Fang et al. 2017). However, the upstream molecular mechanisms controlling the wide range of nitrate response genes remain elusive.

NODULE INCEPTION (NIN) is the first transcription factor identified as essential for nodulation and the symbiotic nitrogen fixation pathway in *Lotus japonicas* (Schauser et al. 1999). NIN-like proteins (NLPs) encoded by other higher plants are proteins having homology to NIN. In total, the *A. thaliana* and rice genomes encode nine and six NLPs, respectively. Rice and *A. thaliana NLPs* are expressed in almost all organs (Chardin et al. 2014). The most conserved region in NLPs is the RWP-RK domain, possibly involved in DNA-binding and dimerization. In rice, amino acid sequences corresponding to the RWP-RK domain have diversified significantly more than in *A. thaliana* (Schauser et al. 2005). But the RWP-RK domain’s activity is unrelated to nitrate signaling, whereas the N-terminus function as a transcriptional activation domain (Konishi and Yanagisawa 2013). NLPs carry another conserved PBI domain, a protein-protein interaction domain, enabling heterodimerization between PB1 domain containing proteins at their C-terminus (Schauser et al. 2005).

In several previous studies in *A. thaliana*, NLPs are identified as nitrate-responsive *cis*-element (NRE)-binding proteins with transcriptional activation roles and function in the nitrate-regulated expression of a number of genes. For example, suppression of *AtNLP6* down-regulates nitrate-inducible genes, including nitrate transporter genes (*NRT1.1* and *NRT2.1*), nitrate assimilation-related enzyme genes (*NIA1*, *NIA2*, *NIR1*, and *UPM1*) and transcription factor genes (a GARP-type transcription factor gene and *LBD39*) (Konishi and Yanagisawa 2013). Overexpression of *AtNLP7* improves plant growth under both nitrogen-poor and nitrogen–rich conditions by coordinately enhancing nitrogen and carbon assimilation (Yu et al. 2016). *AtNLP8* is essential for nitrate-promoted seed germination, reducing abscisic acid levels in a nitrate-dependent manner (Yan et al. 2016). On the other side, the NRE region (a 43 bp-long palindromic sequence) does not exist in genes other than *NIA1* and *NIR1* in *A. thaliana*. In contrast, in maize the NRE region is found to be over-represented in the promoter of genes coding nitrate transporters and ammonium transporters (Liseron-Monfils et al. 2013). In rice, NRE-like sequence comprises two half-sites, 5’-AAG(A/G)GCC-3’ and 5’-GCCCTTT-3’ separated by 10 nucleotides (Schauser et al. 2005).

It has also been demonstrated that no *NLP* gene is significantly induced by nitrate treatment in *A. thaliana* (Konishi and Yanagisawa 2013). The subcellular localizations of NLPs have also been reported in *A. thaliana*. AtNLP8 localizes to nuclei with and without nitrate treatment (Yan et al. 2016) but a rapid accumulation of AtNLP7 in the nucleus is induced by nitrate (Marchive et al. 2013). Furthermore, AtNLP6 is reported to serve as a partially redundant activator with AtNLP7. The teosinte branched1/cycloidea/proliferating cell factor1-20 (TCP20) can interact with AtNLP6&7 to construct TCP20- NLP6&7 heterodimers functioning as the active complexes to control plant responses to nitrate availability (Guan et al. 2017). AtNLPs are also found to be activated post-transnationally. Upon nitrate supply, an inactive form of AtNLPs is converted into an active form (Konishi and Yanagisawa 2013). A uniquely conserved serine (Ser205) is identified in AtNLP7 as an important phosphorylation site for converting to an active form and nitrate-triggered nuclear retention. The subgroup III Ca^2+^-sensor protein kinases (CPK10, CPK30, and CPK32) are master regulators in primary nitrate signaling and phosphorylate the nitrate-responsive domain of AtNLP7 *in vivo* in the presence of nitrate (Liu et al. 2017a).

Nevertheless, little is known about the role of OsNLP family in rice, except that OsNRT1;1 affect the subcellular localization of OsNLP4 (Wang et al. 2018). In the present study, we investigated *OsNLP4* is an important gene on growth and nitrate accumulation under a condition which nitrate was used as the sole nitrogen source. Our results revealed the high mRNA expression level of *OsNLP4* under no supplemental nitrogen condition and down-regulated expression of genes related to nitrate assimilation in the *osnlp4* mutant lines. In addition, our research also suggested that unlike AtNLP7, OsNLP4 did not show nitrate-promoted nucleocytosolic shuttling mechanism. The paddy field data showed *OsNLP4* has a vital effect on rice production. The role and mechanism of OsNLP4-mediated regulation of nitrate-promoted growth were discussed.

## Results

### *OsNLP4* is essential for nitrate-dependent rice growth

To investigate the function of the rice NLP family, we obtained an *OsNLP4* Tos-17 insertion line (*osnlp4-1*) which has a Tos-17 insertion in the second exon (in the front of ^1044^A) in the Nipponbare background. A homozygous mutant line in terms of Tos-17 insertion in *OsNLP4* were established and they were grown under the 2 mM ammonium or 2 mM nitrate conditions for two weeks. When grown on the modified Kimura B solution in which nitrate was used as sole inorganic N source, the mutant line showed growth defects compared with that of wild type (Figure 1A). The *osnlp4-1* plants grown hydroponically on 2 mM nitrate showed shorter shoot length and root length, almost 70% of wild type (Figure 1B, C). In addition, the shoot dry weight and root dry weight were reduced in the *osnlp4-1* by 58% and 53%, respectively (Figure 1D, E). Such reduction in growth was not observed when plants were grown with ammonium (Figure 1B, C, D, E). These results suggested that *OsNLP4* plays an important role in nitrate dependent growth in rice.

**Figure 1.**
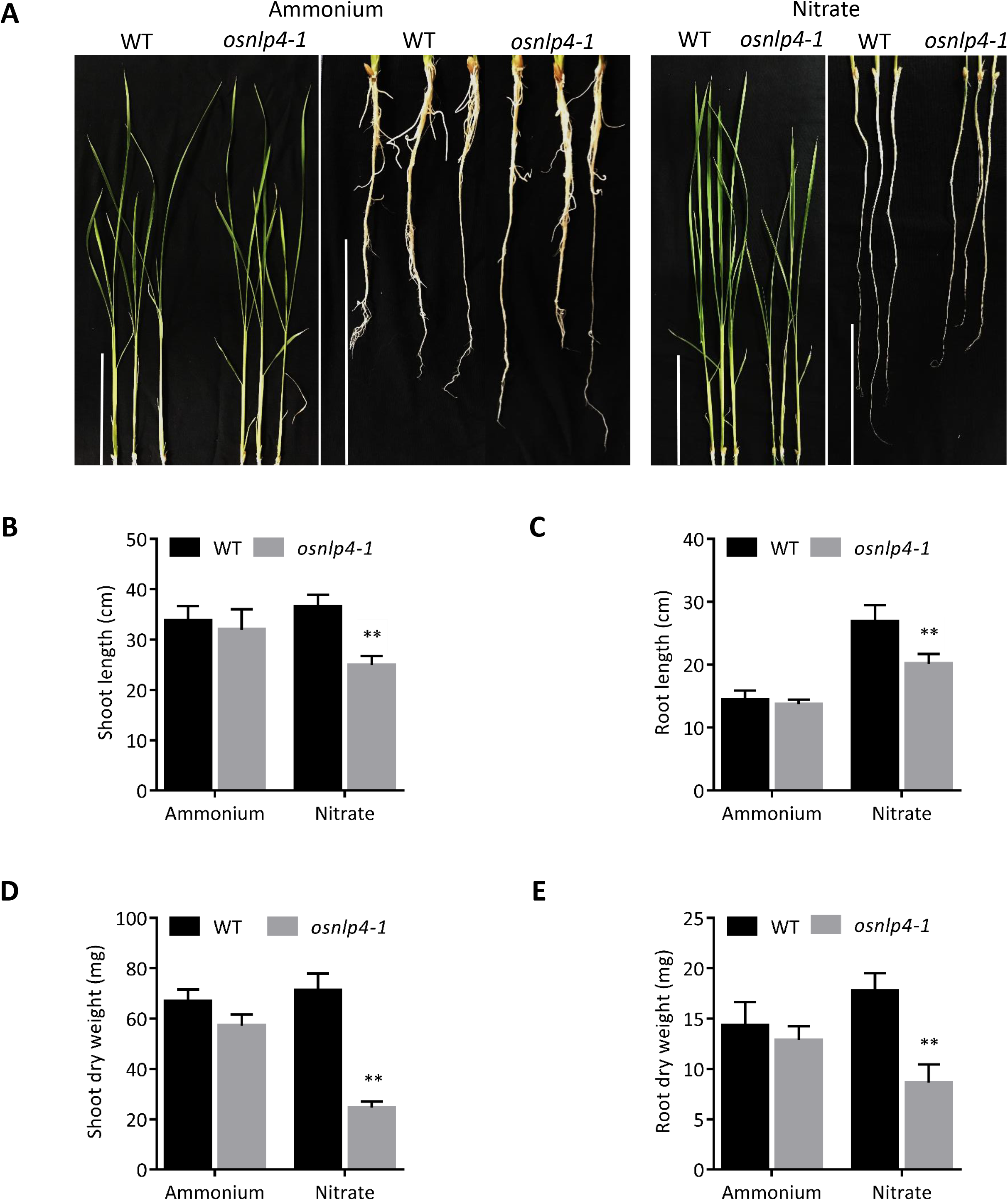
Morphological and physiological phenotype of *OsNLP4* Tos-17 insertion line. **A**, Plants grown in hydroponic culture for 15 days on 2 mM ammonium (NH_4_Cl) or 2 mM nitrate (KNO_3_) modified KimuraB solution with of a density of 8 seeds/L. Scale bars, 10 cm. The shoot length (**B**), root length (**C**), shoot dry weight (**D**) and root dry weight (**E**) of the *osnlp4-1* were compared to wild type for plants grown as in **A**. Error bars represent the standard deviation for 6 plants. Differences are statistically significant (t-test, \*\**p*<0.01).

To further verify if the phenotype of *osnlp4-1* is due to the disruption of the *OsNLP4* gene, we constructed two *OsNLP4* CRISPR/Cas9 lines, *osnlp4-2 and osnlp4-3*. The *osnlp4-2* has 2 bp deletion in the second exon, while *osnlp4-3* has 1 bp insertion in the fourth exon of gene *OsNLP4* (Figure 2A). The *osnlp4-2 and osnlp4-3* reduced shoot length, root length, shoot dry weight and root dry weight only under nitrate condition (Figure 2B, C, D, E). The fact that all the three independently obtained *osnlp4* mutants exhibited impaired growth under nitrate condition, but not under ammonium condition, established that mutations in *OsNLP4* are responsible for these phenotypes.

**Figure 2.**
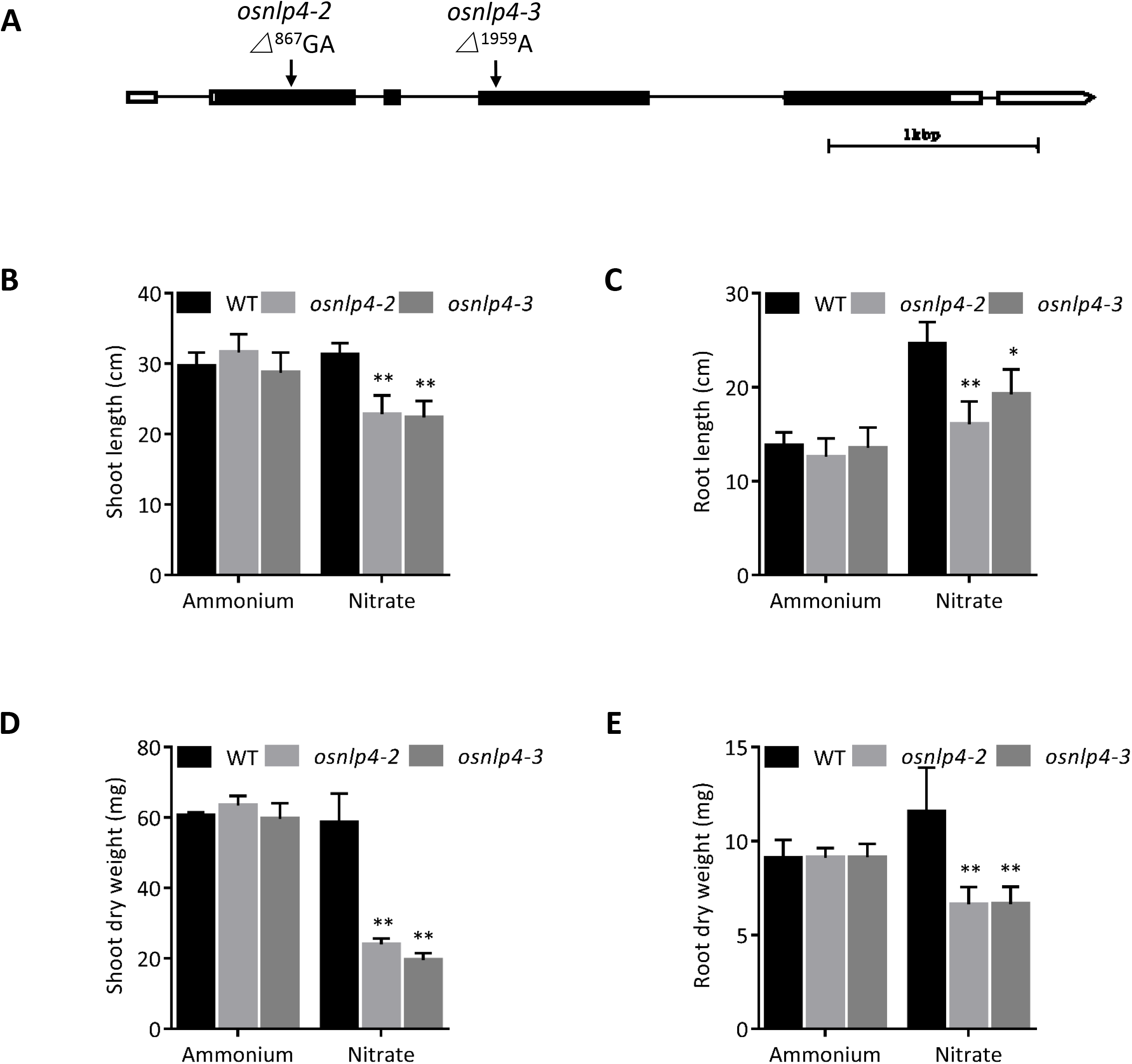
Morphological and physiological phenotype of *OsNLP4* CRISPR/Cas9 lines. **A**, The CRISPR/Cas9 lines of *OsNLP4*. The *osnlp4-2* has 2 bp deletion in the second exon, while *osnlp4-3* has 1 bp insertion in the fourth exon. Lines represent introns, whereas boxes represent exons. The shoot length (B), root length (C), shoot dry weight (D) and root dry weight (E) of *osnlp4-2* and *osnlp4-3* compared to wild type for plants grown in 2 mM KNO_3_. Error bars represent the standard deviation for plants. Differences are statistically significant (Dunnett’s test, \**p*<0.05, \*\**p*<0.01).

### *OsNLP4* mRNA accumulation was increased significantly under nitrogen free condition

All six members of rice NLP gene family (*OsNLPs*) encompassed RWP-RK domain and PBI domain. *OsNLPs* can be classified into three clades (*OsNLP1* and *OsNLP4*, *OsNLP2* and *OsNLP5*, *OsNLP3* and *OsNLP6*) (Supplemental Figure S1).

Here we determined mRNA accumulation of *OsNLP* genes in rice. We performed RNA-seq analyses using RNA extracted from roots of 15-day wild type plants grown under 2 mM KNO_3_ (nitrate), 2 mM NH_4_Cl (ammonium), 1 mM KNO_3_ + 1 mM NH_4_Cl (normal) or 2 mM KCl (nitrogen free). Compared with the normal condition, the mRNA accumulations of a number of genes were up-regulated or down-regulated under the other three conditions. Under the nitrogen free condition, 2062 genes were up-regulated specifically (Figure 3A). *OsNLPs* were among these genes except *OsNLP3* and *OsNLP6*. Especially, *OsNLP1* and *OsNLP4* had significantly higher mRNA expression levels (Figure 3B). Expression of *OsNLPs* was further analyzed by quantitative real-time PCR (qRT-PCR) to detect their mRNA expression levels under the nitrate and the nitrogen free conditions. The result was consistent with RNA-seq data. *OsNLP4* was increased significantly only under the nitrogen free condition in both shoots and roots (Figure 3C). This implies possible involvement of OsNLP4 under the nitrogen free condition, but the *osnlp4* mutant lines did not show apparent difference in physiological phenotype with wild type under the nitrogen free condition (Supplemental Figure S2A, B, C, D, E).

**Figure 3.**
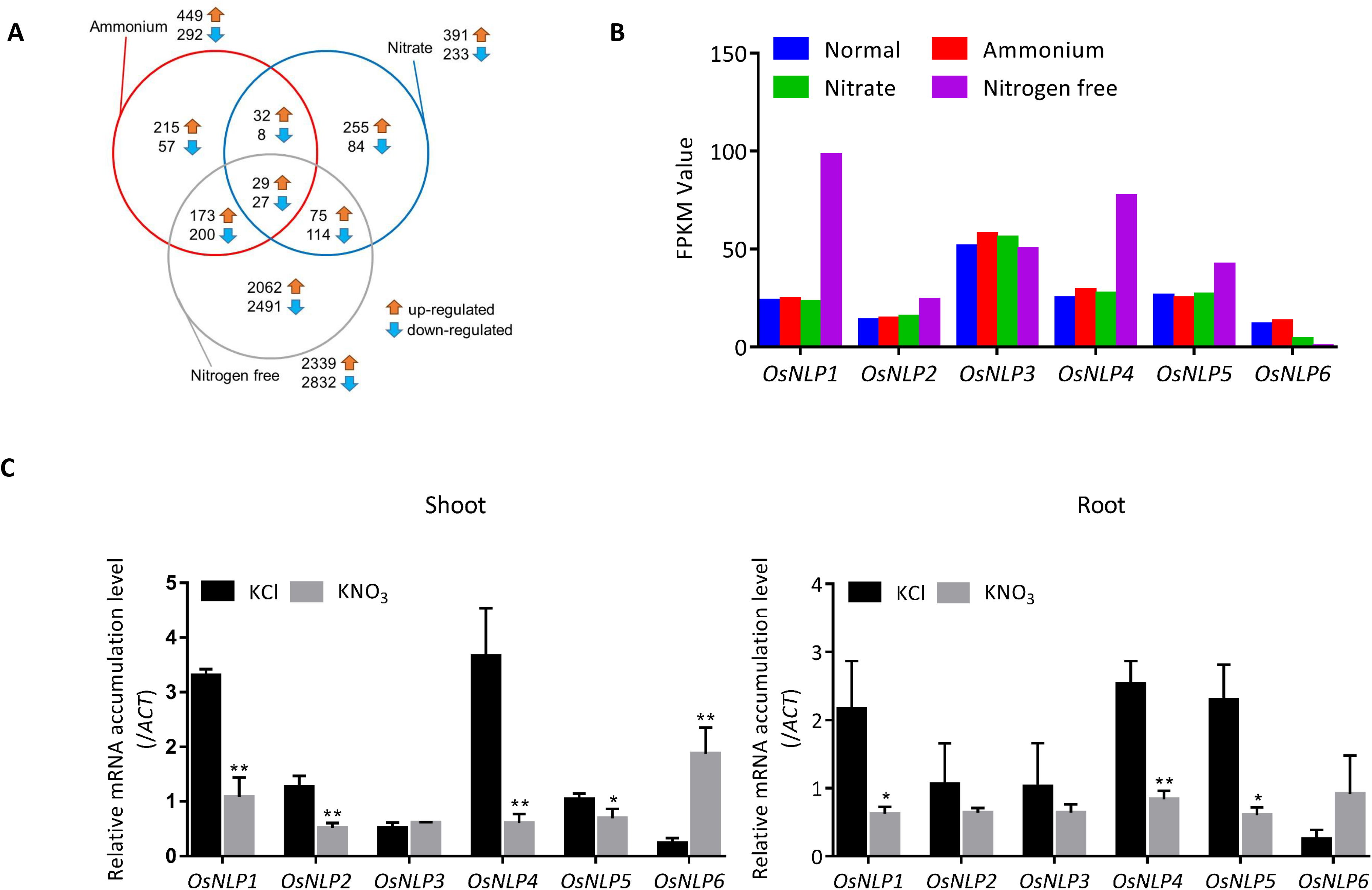
The mRNA expression levels of *OsNLPs* in wild type under a range of nitrogen conditions. Total RNA was extracted from 15-day-old plants and was subjected to RNA-seq and qRT-PCR. **A**, Crossing pin charts were used to present the comparison and overlaps between up-regulated and down-regulated genes defined under different nitrogen conditions (Nitrate, 2 mM KNO_3_; Ammonium, 2 mM NH_4_Cl; Nitrogen free, 2 mM KCl) compared with normal condition (1 mM KNO_3_ + 1 mM NH_4_Cl) according to RNA-seq data. **B**, RNA-seq data showed transcript accumulations of six *OsNLP* family members in wild type under four different nitrogen conditions. **C**, Relative expression of *OsNLP*s was analyzed by qRT-PCR and normalized to the expression of *ACT*. Each gene had three biological replicates and the assays were repeated two times. Differences are statistically significant (t-test, *p*<0.05, \*\**p*<0.01).

Taken together, our results suggested that the accumulation of *OsNLP4* transcript in rice is induced by nitrogen starvation.

### Suppression of *OsNLP4* broadly down-regulates the expression of genes involved in nitrate assimilation

To further determine the origin of the mutant’s N-starved like phenotypes and assess the role of OsNLP4 in nitrate signaling, RNA-seq experiment was performed using RNA extracted from wild type and the *osnlp4-1* roots grown for 15 days on the nitrate condition. RNA-seq data showed there were 580 genes up-regulated and 622 genes down-regulated for the *osnlp4-1* compared with wild type. Most genes are related to catalytic activity, transporter activity and binding in both parts (Supplemental Figure S3A, B). Genes related to nitrate assimilation (*NIA*, nitrate reductase gene; *NIR*, nitrite reductase gene) were among those down-regulated genes (Supplemental Figure S3C). In addition, the expression levels of *IRT1* (encoding iron transporter) and *MOT1* (encoding molybdenum transporter) were also impaired in the *osnlp4-1* (Supplemental Figure S3C).

Expression of some selective genes was further analyzed by qRT-PCR to examine their mRNA expression levels. Result indicated that in roots, mRNA accumulations of *NIA1* and *NIR1* were declined in the *osnlp4* mutants compared to the wild type (Figure 4). On the other hand, genes related to nitrate transport, i.e., *NRT2.1*, high-affinity nitrate transporter; *NRT1.5a*, root-to-shoot transporter, had similar expression levels between wild type and the *osnlp4* mutants (Figure 4). In shoots, the accumulations of *NIA2* and *NRT2.1* transcripts were increased in the *osnlp4* mutants while *NIA1* was reduced (Figure 4). These results indicated that mRNA accumulation of genes related to nitrate assimilation but not those related to nitrate transport are impaired in the *osnlp4* mutants.

**Figure 4.**
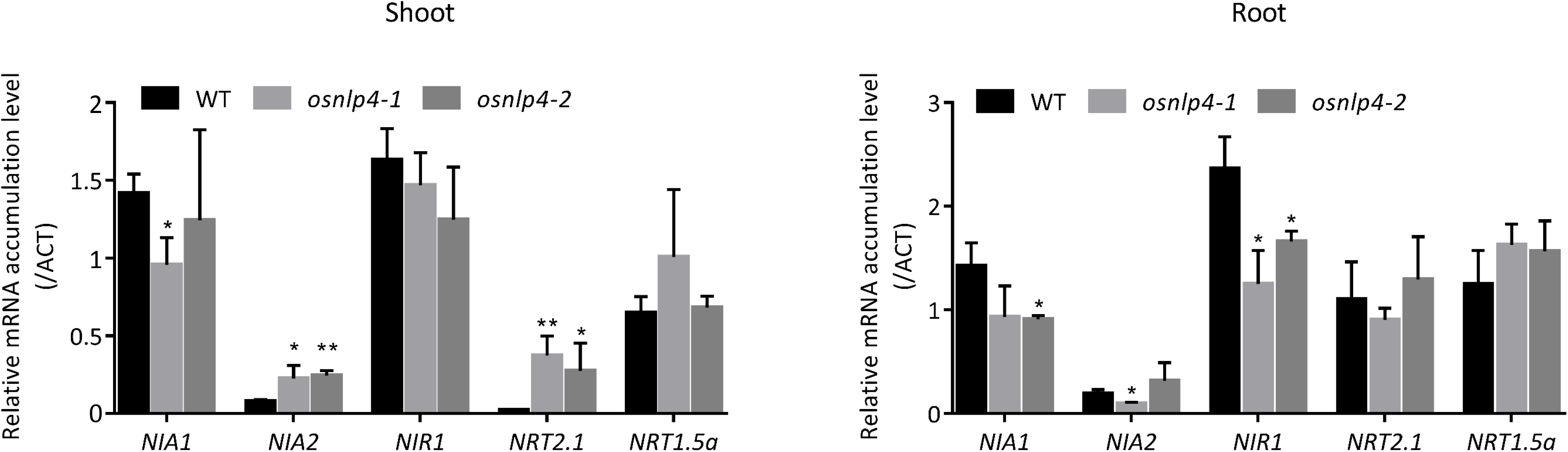
The mRNA expression levels of nitrate reductase genes and nitrate transporter genes in the *osnlp4* mutants under nitrate condition. The steady-state transcript amounts for the nitrate reductase genes *NIA1* and *NIA2*, nitrite reductase gene *NIR1*, high-affinity nitrate transporter genes *NRT2.1* and root-to-shoot nitrate transporter gene *NRT1.5a* were determined by qRT-PCR. The results given are a percentage of the level of mRNA for the *ACT* gene. Error bars, s .d, n=3 biological replicates (Dunnett’s test, \**p*<0.05, \*\**p*<0.01).

Overall, our analysis suggest that OsNLP4 plays a role in nitrate regulation of gene expression including genes involved in nitrate metabolism.

### OsNLP4 is involved in nitrate uptake and assimilation

To examine physiological defects of *osnlp4* mutants, we determined the nitrate absorption rate of wild type and the *osnlp4* mutants under continuous nitrate and one-day nitrate starvation conditions. It was found that nitrate uptake rate of the *osnlp4* mutants were lower than that of the wild type in both conditions (Figure 5A, B). In particular, after one-day nitrate starvation treatment, wild type showed high nitrate absorption rate at almost 1.5 NO_3_^−^ μmol/root fresh weight (g)/h while the *osnlp4* mutants displayed less than half of that of wild type (Figure 5B).

**Figure 5.**
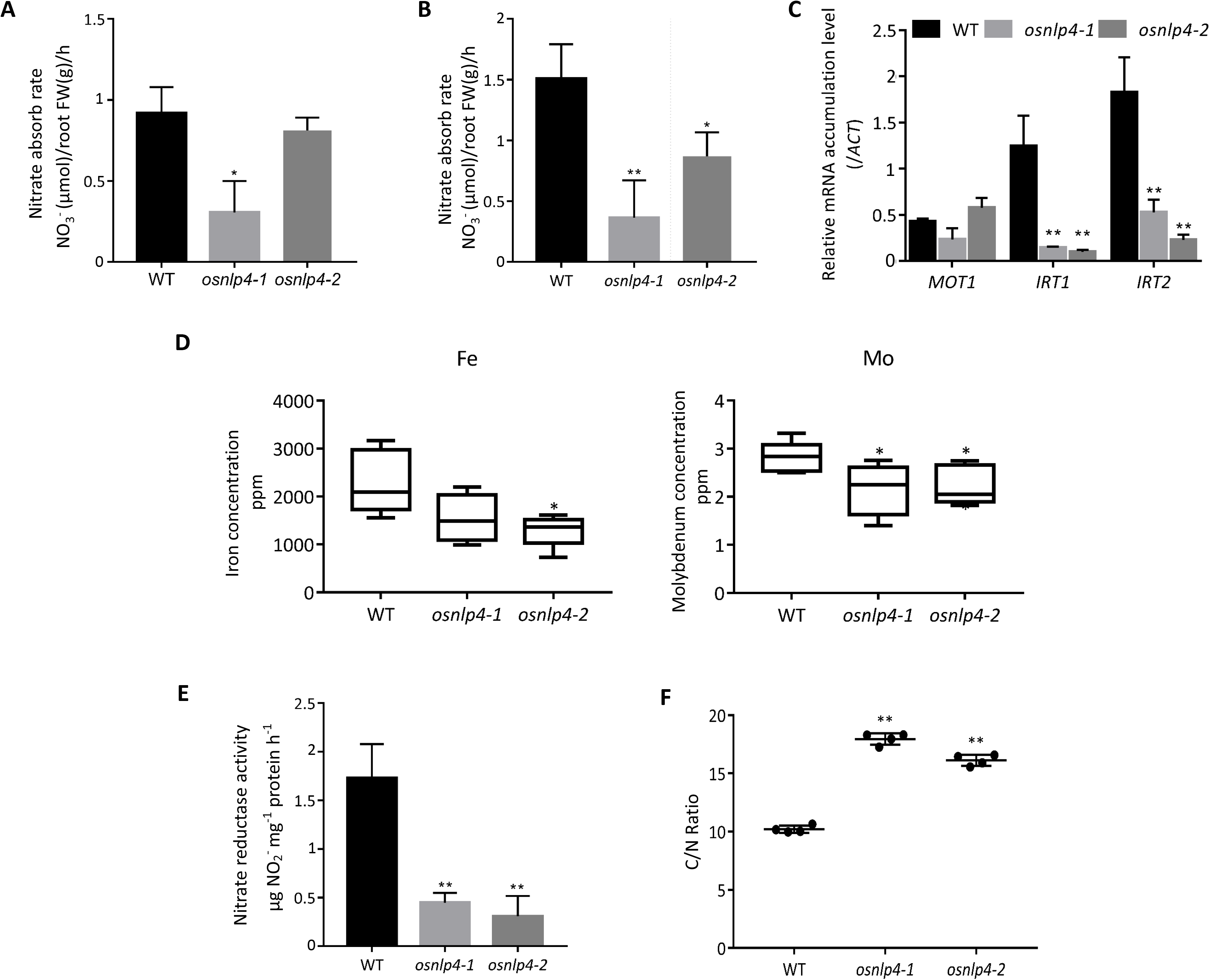
The *osnlp4* mutants impair nitrate uptake rate and nitrate assimilation. **A**, Nitrate absorb rate of WT and *osnlp4* mutants under continuous nitrate condition. **B**, The14-day old plants grown on 2 mM KNO_3_ were transferred to 2 mM KCl. After one day, removed plants to 2 mM KNO_3_ again and detected nitrate absorb rate. **C**, The mRNA expression levels of molybdenum transporter gene *MOT1* and iron transporter genes *IRT1*, *IRT2* in root were determined by qRT-PCR. The mRNA accumulations were normalized by that of *ACT.* **D**, Iron and molybdenum concentrations in root. **E**, Nitrate reductase activity in root. **F**, The carbon to nitrogen ratio of the *osnlp4* mutants. Error bars, s .d, n=3 biological replicates (Dunnett’s test, \**p*<0.05, \*\**p*<0.01).

In the aforementioned RNA-seq result, *MOT1* and *IRT1* mRNA accumulations were reduced in the *osnlp4-1* (Supplemental Figure S3C). Molybdenum and iron have been reported to be essential for the activity of nitrate reductase through covalently bounding to specific domains of the enzyme (Liu et al., 2017b, Ford et al., 2016). We checked the mRNA expression levels of *MOT1*, *IRT1* and *IRT2* in root via qRT-PCR. The expressions of *IRT1* and *IRT2* were strongly impaired in the *osnlp4* mutants (Figure 5C). And the total content of iron and molybdenum was declined in the *osnlp4* mutants (Figure 5D). It is possible that NR activity is affected by OsNLP4. We compared the nitrate reductase (NR) activity of the *osnlp4* mutants with WT grown under nitrate condition and discovered that NR activity was lower in the *osnlp4* mutants, almost only 30% of WT, establishing that OsNLP4 is required to maintain NR activity (Figure 5E).

We next analyzed shoot nitrogen content to verify that OsNLP4 is important for rice to assimilate nitrate by CN coder. The C/N ratio (carbon-to-nitrogen ratio) in shoot was nearly two-fold higher in the *osnlp4* mutants under nitrate condition which means the nitrogen limitation existed in the *osnlp4* mutant lines (Figure 5F).

These results collectively suggested that OsNLP4 is indispensable for the expression of genes involved in then nitrate transduction pathway both in nitrate uptake and assimilation. OsNLP4 also affects nitrate use efficiency, probably through the regulation of nitrate uptake and assimilation.

### OsNLP4 is localized in nucleus

In *A. thaliana*, AtNLP6 and AtNLP7 are retained in the nucleus in the presence of nitrate, but AtNLP8 are not (Yan et al., 2016, Yu et al., 2016, Guan et al., 2017) and these changes in subcellular localization is implied as an important step for nitrate dependent gene expression, we tested whether the subcellular localization of OsNLP4 is under the control of nitrate conditions. Rice protoplasts were transiently transformed with 35S::*OsNLP4*::*GFP*. After 18h incubation with or without nitrate, OsNLP4-GFP signal was detected mainly in the nucleus with minor signals and these pattern of subcellular localization does not seem to be affected by the nitrate condition (Figure 6). These results indicated that OsNLP4 is localized to nucleus irrespective of nitrate conditions. This is in contrast to the case of AtNLP7, implying that OsNLP4 in rice is regulated by a different mechanism.

**Figure 6.**
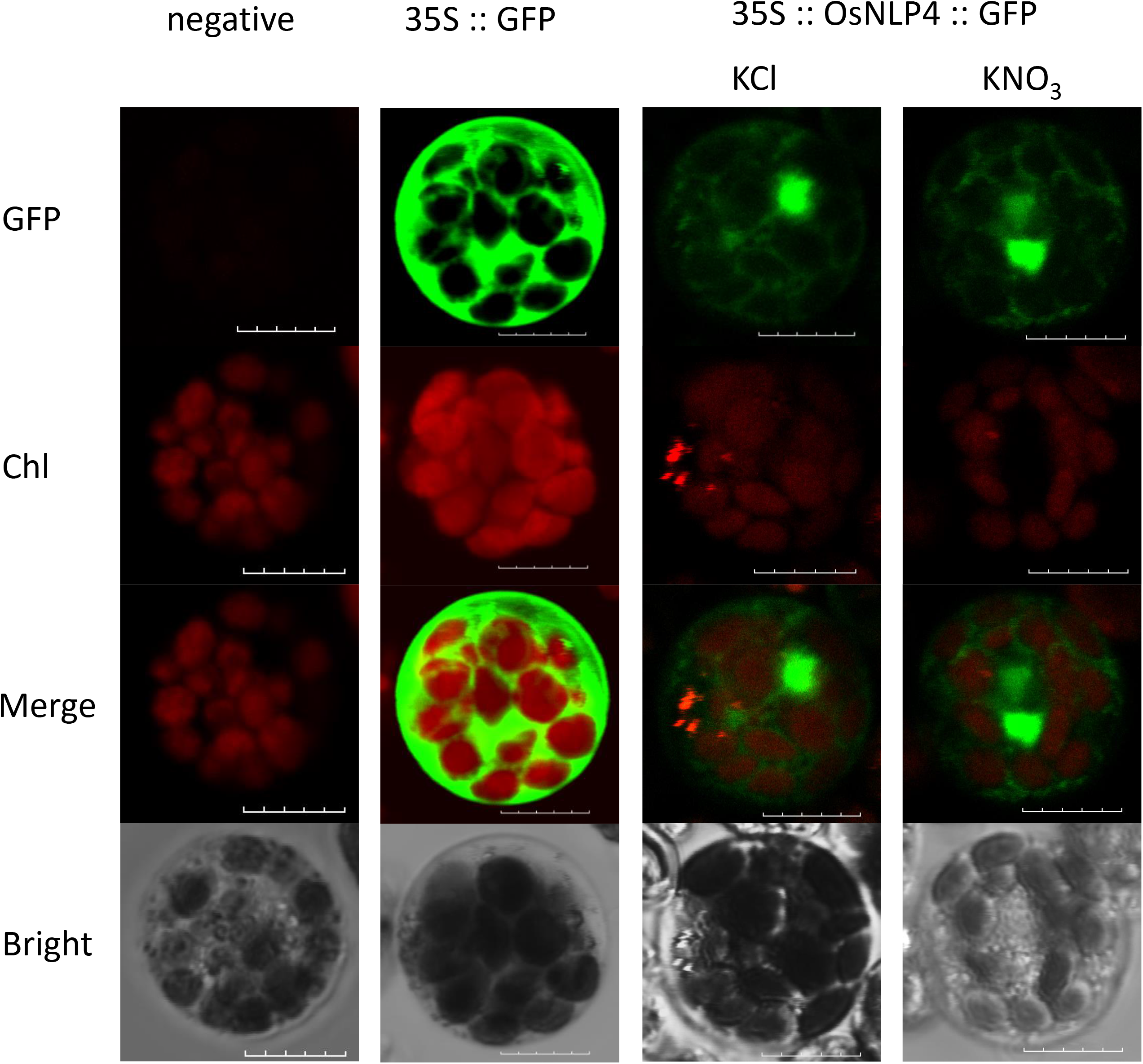
Subcellular localization of OsNLP4 :: GFP in the KCl- or KNO_3_- treated rice protoplasts. The pMDC83 vector was used as a negative control and 35S :: GFP was used as a positive control of GFP signal. After PEG-mediated transformation, added KCl or KNO_3_ to the protoplast incubation medium at 2 mM final concentration. Chl : chloroplasts autofluorescence. A bar indicates 10 μm. Images are representative of 10 protoplasts.

### The *osnlp4-1* line has low rice yield in paddy field

Rice is the main cereal crop in substantial portions of the world and nitrogen fertilizers are frequently used with the aim to achieve high yields. In order to check agronomic traits of our *OsNLP4* Tos-17 insertion line, we planted WT (Nipponbare) and the *osnlp4-1* in nitrogen-supply (+N) and no-nitrogen supply(−N) plots in the Kashimadai paddy field of Tohoku University (Osaki-city, Japan) in 2017, 2018, and 2019 (Figure 7A, B, C). Honestly, these three years data are not consistent with each other very well because of many uncontrollable factors, like temperature, typhoon etc. Although there was no significant difference of tiller number between WT and the *osnlp4-1*, the *osnlp4-1* line showed the trend of decreased productive panicle weight and straw weight (Figure 7A, B, C). Significant difference on grain yield was also observed between WT and the *osnlp4-1* line in the no-nitrogen supply plot (Figure 7A, C). It is possible that residual nitrogen existed in no-nitrogen supply plot. These results indicated the importance of OsNLP4 in yields in paddy fields. We believe that the study of *OsNLP4* has a potential to increase rice yield in paddy fields.

**Figure 7.**
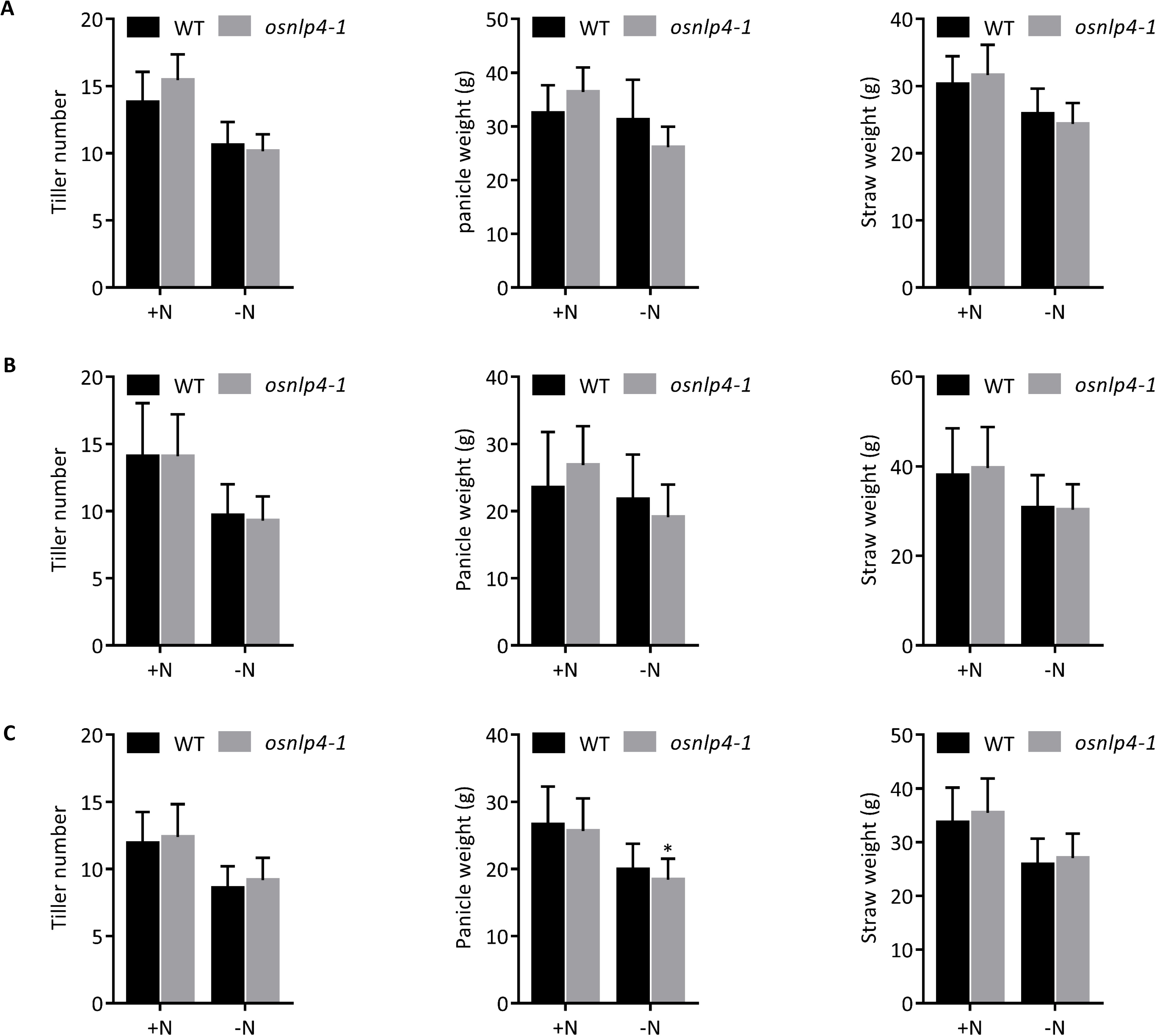
Agronomic traits of the *osnlp4* Tos-17 insertion line in paddy field. Tiller number, panicle weight and straw weight per plant were determined for the *osnlp4-1* in comparison to WT in a field trial conducted in Sendai, Japan in 2017 (A), 2018 (B), 2019 (C) (+ N, Nitrogen-supply paddy field; −N, No-Nitrogen supply paddy field). Error bars represent s .d, n = 20, 40, 68 (t-test, \**p*<0.05, \*\**p*<0.01).

## Discussion

### Rice accumulates *OsNLP4* at RNA level under no nitrogen condition

Most of the genes involved in nitrate signaling are induced by nitrate itself, although its mode of action is poorly understood (Hoff et al. 1994; Krouk et al. 2010a; Gutierrez 2012; Krapp et al. 2014; Konishi and Yanagisawa 2014). Recently, NLPs had proved to work as key transcription factors responsible for nitrate-inducible expression of a number of genes including nitrate transporter, nitrate reductase, nitrite transporter and nitrite reductase in response to nitrate in *A. thaliana* (Yu et al. 2016; Castaings et al. 2009; Yanagisawa 2014). AtNLPs can directly bind to the NRE domain within some nitrate-induced genes probably through post-translational activation (Konishi and Yanagisawa 2013). Our present study identified *OsNLP4* as an important gene for regulating growth under nitrate condition in rice. The *osnlp4* mutants displayed prominent nutrient deficiency symptom when nitrate was used as sole nitrogen source (Figure 1A, B, C, D, E & Figure 2B, C, D, E). Notably, qRT-PCR results showed that mRNA expression level of *OsNLP4* was increased significantly under no nitrogen condition (Figure 3B, C). In *A. thaliana*, no *AtNLP* was induced by N source or N starvation (Konishi and Yanagisawa 2013), but *AtNLP8* and *AtNLP9* were highly induced in imbibed seeds (Yan et al. 2016). One possibility is that OsNLP4 is inactive when no nitrate is supplied. We speculate that rice accumulates a high level of OsNLP4 in order to react to nitrate quickly, converting the inactive form of OsNLP4 into active form when nitrate signaling is sensed.

Among the OsNLP family, in addition to OsNLP4, it seemed like that rice also accumulate OsNLP1 and OsNLP2 especially in shoots and roots under nitrate starvation while OsNLP6 was the only repressed one (Figure 3B, C). OsNLP6 has a conserved N-terminal part of the RWP-RK domain, but lacks the downstream half (Schauser et al. 2005). This protein might have lost putative ancestral DNA-binding function and have acquired a divergent function. Our results support this hypothesis that NLPs for a specific functional role is different among plant species.

### OsNLP4 triggers a number of nitrate specific responses

We found 622 down-regulated genes in the *osnlp4* mutants dependent of nitrate application, among which most were genes related to catalytic activity (Supplemental Figure S3B). Transcript accumulations of *NIA1*, *NIA2*, *NIR1*, *IRT1*, *MOT1*, and *NAS1* in the *osnlp4* mutants were lower than WT in root (Supplemental Figure S3C & Figure 5C). Nitrate reductase requires iron and molybdenum to consist co-factors to catalyze the reduction of nitrate-to-nitrite in cytosol. And the reduction needs NAD(P)H as energy (Hoff et al. 1994). As expected, the NR activity was impaired in the *osnlp4* mutants (Figure 5E). Reduced NR activity further leaded to low total nitrogen content and negatively affected growth in *osnlp4* mutants under nitrate-sufficient condition (Figure 5F & Figure 1A). On the other hand, no obvious difference was detected of *NRT2.1* and *NRT1.5a* mRNA levels between WT and the *osnlp4* mutants in roots. In shoots, the mRNA expression levels of *NIA2* and *NRT2.1* were even higher than WT in the *osnlp4* mutants (Figure 4). As reported, both nitrate assimilation and nitrate transport were impaired in *atnlp7* mutants (Castaings et al. 2009). On the basis of our results, we propose that OsNLP4 mainly regulates the expression of nitrate assimilation genes but not nitrate transporter genes in root, acting as a nitrate sensor to trigger secondary responses. It is possible that OsNLP4 displays different requirements for NRE-like motif recognition and has different functions in shoots and roots.

### OsNLP4 does not have nitrate-promoted nucleocytosolic shuttling mechanism

The localization or retention of AtNLP7 in nuclei is controlled by nitrate signaling and the phosphorylation of a conserved serine residue is essential for the nitrate-simulated rapid nuclear translocation (Marchive et al. 2013; Liu et al. 2017a). However, OsNLP4-GFP was localized to the nucleus independent of nitrate and did not show nitrate-promoted nucleocytosolic shuttling mechanism in our experimental condition (Figure 6). Wang et al. (2018) demonstrated possible shuffling mechanisms of OsNLP3 and OsNLP4 in rice upon exposure to 10 mM nitrate. In our experiments we used 2 mM nitrate which is the condition required to observe growth phenotype of *osnlp4* mutants. It is possible that shuffling of OsNLP4 between cytoplasm and nucleus does not occur at 2 mM but occurs at 10 mM and the role of OsNLP4 maybe different between 2 mM and 10 mM nitrate conditions (Wang et al. 2018). It is known that except nitrate multiple signals coordinately interact with each other through NLPs in plant to modulate downstream events, such as Ca^2+^ signaling, phytohormones and drought (Guan 2017; Almeida et al. 2017; Liu et al. 2017a). It is possible that multiple mechanisms that activate NLPs functions exist in different plant species.

### A schematic model of proposed OsNLP4 action in rice

We propose a working model to illustrate how OsNLP4 might react to nitrate signaling to regulate the nitrate assimilation pathway in rice. Nitrate is perceived as signal by some unknown sensors, then transmitted via a signal transduction pathway towards OsNLP4. OsNLP4 controls the expression of a wide range of genes related to nitrate assimilation that have an impact on NR activity and other developmental processes (Figure 8). In addition, the expression of nitrate-assimilation related genes is also regulated by various environmental factors. Probably, crosstalk may occur between OsNLPs and other transcription factors to coordinate the nitrate assimilation pathway.

**Figure 8.**
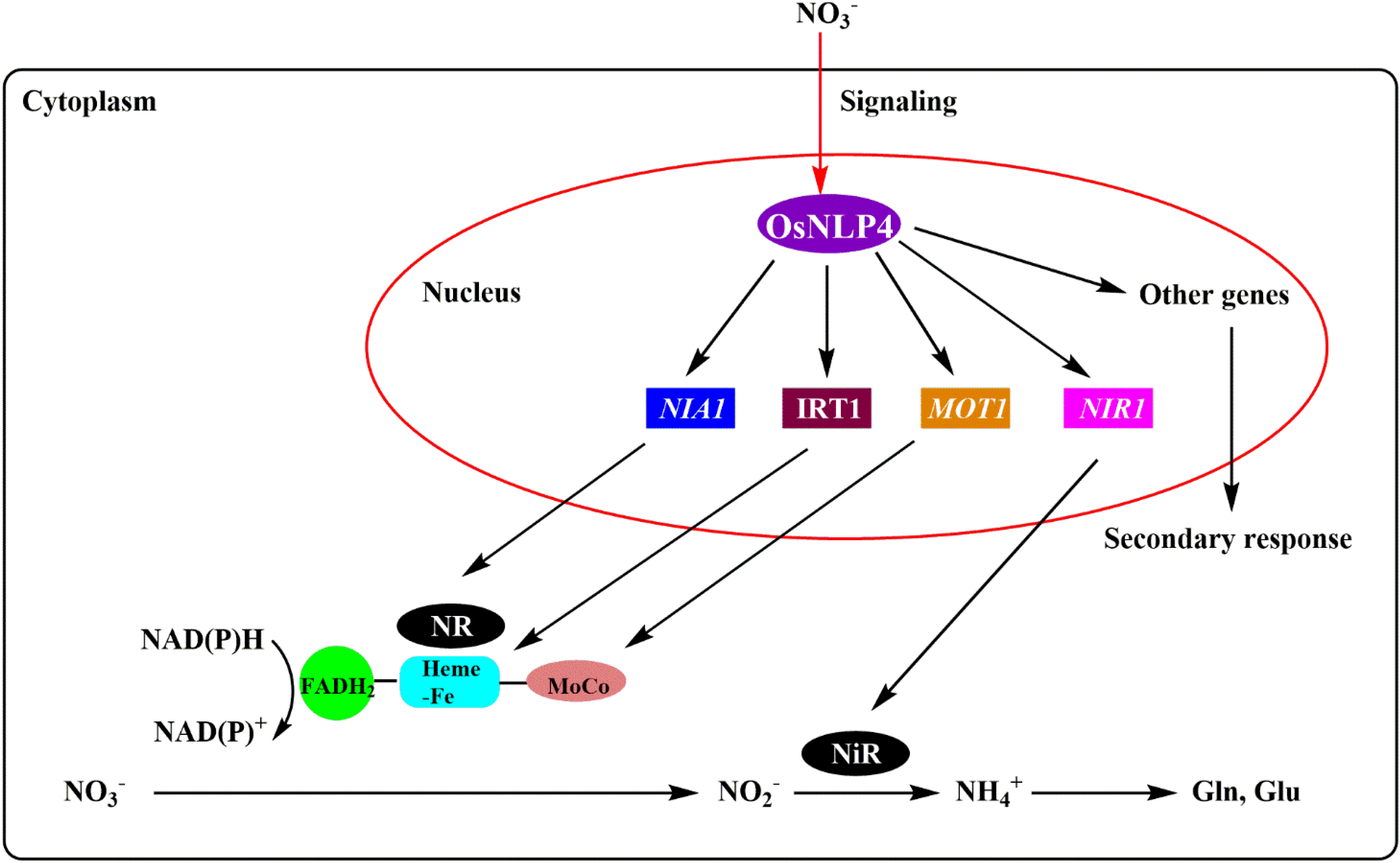
A schematic model of proposed OsNLP4-regulated nitrate assimilation in response to nitrate provision in rice. Nitrate is perceived as signal and transmitted via unknown pathway towards transcription factor OsNLP4. Then OsNLP4 promotes the expression of nitrate assimilation-related genes and other unknown regulatory genes. High active nitrate reductase catalyzes nitrate reduction to trigger nitrate specific growth responses.

Recently, it is reported that in *A. thaliana* NLPs also take a role in developmentally programmed processes independent of nitrate. AtNLP7 can regulate root cap cell release likely through regulation of *CELLULASE5* (Karve and Iyer-Pascuzzi 2018). So further studies are clearly required to understand how OsNLP4 reacts to nitrate signaling with changing from inactive form to active one and the functions of other OsNLPs in rice. NLPs might take a part in abundant developmental and metabolic processes as a transcription factor in plant. Identification and characterization of NLP family probably provide clues for understanding nitrogen or other metabolic pathway.

### *OsNLP4* have potential effect on sustainable agriculture

Furthermore, nitrogen use efficiency (NUE) is an important indicator for the development of sustainable agriculture (Xu et al. 2012). In order to increase grain yield, farmers use high amount of chemical fertilizers, especially nitrogen fertilizers, but with considerable negative impacts on the environment (Han et al. 2015). Our paddy field data indicated that the *osnlp4* impaired grain yield (Figure 7). The elevated gene expression in *OsNLP4* might enhance NUE in rice paddy field. Comprehension of the function of *OsNLP4* involved in nitrate signaling may increase crop growth and productivity under nitrogen constraints.

## Materials and Methods

### Plant materials and growth conditions

*Oryza sativa* L. cv. Nipponbare was used as the wild type. The *OsNLP4* Tos-17 insertion line (*osnlp4-1*), which has an insertion in second exon, was obtained from the collection of Tos-17 mutant panel by National Institute of Agro Biological Resources, Tsukuba, Japan. Homozygous Tos-17 line was identified using *OsNLP4* gene-specific primers and Tos-17 left-border primer. To generate *OsNLP4* CRISPR/Cas9 lines (*osnlp4-2* and *osnlp4-3*), sgRNAs designed on CRISPR-P (http://crispr.hzau.edu.cn/CRISPR2/) were annealed into pU6gRNA vector (Shan et al. 2013) and subsequently cloned into the modified binary vector pZDgRNA_Cas9 ver.2_HPT (Endo et al. 2015). All constructs were sequence verified and transformed into Nipponbar*e* using the *Agrobacterium*-mediated rice transformation method. The *osnlp4-2* line has a ^867^GA deletion in second exon and the *osnlp4-3* line has a ^1959^A insertion in fourth exon. The primers used are listed in the Supplementary Table 1.

Wild type and the *osnlp4* lines were surface sterilized with 10% (v/v) bleach for 30 min and then rinsed thoroughly with deionized water. The sterilized seeds were germinated on plastic plate containing moist filter paper for 5 days. Uniform seedlings were selected and then transferred to a tank containing 3 L modified Kimura B solutions (Uraguchi et al. 2009)supplemented with one of the following as the sole N source: 2 mM KNO_3_; 2 mM NH_4_Cl; 1 mM KNO_3_ + 1 mM NH_4_Cl and 2 mM KCl. All the plants were grown in a chamber with a 16-h-light/ 8-h-dark photoperiod, and the temperature was controlled at 30 °C. The solution was refreshed every 7 days. Samples were harvested at 15 days for phenotypic analysis, RNA isolation and protein extraction. For nitrate uptake speed analysis, 14-day old plants grown on 2 mM KNO_3_ were transferred to 2 mM KCl and removed plants to 2 mM KNO_3_ again after one day. For the preparation of rice protoplasts, Nipponbare were grown on 1/2MS medium at 30 °C for 10 days under continuous light.

### RNA isolation and Quantitative real-time PCR

Total RNA was extracted from shoots and roots using NucleoSpin RNA Kit (MACHEREY-NAGEL). 500 ng total RNA was reverse-transcribed using PrimeScript RT Master Mix (Perfect Real Time) (TaKaRa). Quantitative real-time PCR was performed using SYBR Premix Ex Taq II (Tli RNaseH Plus, TaKaRa) and monitored with the Thermal Cycler Dice Real Time System II (TaKaRa). The relative gene expression was normalized to the expression of *OsACT*. Triplicate biological replicates were analyzed. The primers used for qRT-PCR are listed in Supplementary Table 2.

### RNA-seq analyses

Total RNA was used for preparing the library with BGI (https://www.bgi.com/us/) (twofold biological samples). Purification of the Poly-A mRNA was performed using TruSeq RNA Library Preparation (Illumina, https://www.illumina.com/) and paired-end sequencing identified 100-bp sequences at both ends of the cDNA fragments. Fastq files, downloaded from the main facility, were used for data analysis. Reads in the fastq file were mapped to the rice genome, *Oryza sativa* Japonica-IRGSP 1.0.23 (Ensembl Plants http://plants.ensembl.org/index.html) using TopHat2.0.9 (http://ccb.jhu.edu/software/tophat/index.shtml) Finally, Cufflinks2.1.1 (http://cole-trapnell-lab.github.io/cufflinks/) was used to determine differential gene expression and obtain FPKM value.

### Nitrate uptake rate assay

Three 15-day-old seedlings were used for a measurement of nitrate uptake. Three plants were transferred to a 50 ml tube containing 50 ml Kimura B solution with 2 mM KNO_3_ and their roots were rinsed thoroughly with distilled H_2_O. After 3 h and 6 h, sucked 500 μl solution from the tube to detect the concentrations of remaining nitrate using High Performance Capillary Electrophoresis System (Agilent Technologies). Then measured the root fresh weights and calculated the nitrate uptake speeds (NO_3_^−^ (μmol)/root fresh weight (g)/h). Three biological replicates of WT and *osnlp4* mutant lines were analyzed for each treatment.

### Measurement of C/N ratio

The total nitrogen content was investigated using CN coder Vario MAX cube (Elementar). The 15-day shoot samples were dried in a 70 °C oven for 3 days. Then removed samples into assorted metal containers. The machine burned samples and detected total contents of carbon and nitrogen automatically.

### Analysis of NR activity

The nitrate reductase activity was measured by the modified method described by Maeda et al. and Pinto et al. (Pinto et al. 2014). The roots and shoots (1.0 g fresh weight) were separated and ground in a chilled mortar containing 4 ml extraction buffer (50 mM HEPES-KOH pH 8.0, 5 mM EDTA, 10 mM β-mercaptoethanol, 1 mM DTT, 0.5 mM PMSF, 5% (w/v) PVPP). The resulting homogenate was then centrifuged at 13,000 rpm for 30 min at 4 °C. Samples of the supernatant were used for the determination of protein content (NanoDrop ND-1000) and the assays of *in vitro* NR activity (300 μl). The reaction mixture (450 μl) of the latter consisted of 50 mM HEPES-KOH pH 8.0, 1 mM DTT, 2 mM KNO_3_. After incubation at 30 °C for 1 min, the reaction was started by adding 150 μl 0.6 mM NADH. After incubation at 30 °C for 1h, the reaction was ended by the addition of 100 μl cold water. After 1h the NO_2_^−^ produced was colorimetrically measured at 543 nm (SHIMADZU UV-1850) after the addition of 1% (w/v) sulphanilamide dissolved in 2 M HCl and 0.02% (w/v) N-1-naphthylamine. NR activity was expressed as μg NO_2_^−^ mg^−1^ protein h^−1^.

### Protoplast transient assay

To understand the subcellular localization of OsNLP4, p35S-OsNLP4-GFP and p35S-GFP modified from pMDC83 vector (Curtis and Grossniklaus 2003) were transformed into rice protoplasts. The rice protoplast preparation and transfection followed preciously described procedures with some modifications (Bart et al. 2006). Rice protoplasts were isolated from stem of rice seedlings after sowing for 10 days. Briefly, 100 μl of protoplast suspension was transfected with DNA for various constructs using PEG-mediated method. After transfection, cells were cultured with 0.25 ml inoculation solution containing 2 mM KCl or KNO_3_ for 18 h at 25 °C. Cells were collected, then GFP signal in the cells was analyzed using a confocal laser scanning microscope (OLYMPUS FV10-MCPSU).

### Rice plant sampling and analysis

To measure the rice grain yield, WT (Nipponbare) and the *osnlp4-1* line were planted in nitrogen-supply area (+N) and no-nitrogen supply area (−N) of Kashimadai paddy field of Tohoku University (Osaki-city, Japan) in May, 2017/2018/2019. Tree frames (4 lines × 4 lines) were harvested in each area in October, 2017/2018/2019. In the no-nitrogen supply paddy field, phosphorus and potassium were supplied as normal. The panicle weight, straw weight and tiller number were measured from 20 lines sampled from the three harvested frames.

## Accession Numbers

Sequence data from this article can be found in the GenBank/EMBL data libraries under accession numbers: AP014959 (*OsNLP1*), AP014960 (*OsNLP2*), AP014957 (*OsNLP3*), AP014965 (*OsNLP4*), AP014967 (*OsNLP5*), AP014958 (*OsNLP6*).

## Acknowledgments

The authors acknowledge the China Scholarship Council (CSC), Japan Society for the Promotion of Science (JSPS, Nos. 18H05490 and 19H05637) supporting this work.

## Supplemental Data

**Supplemental Figure S1.**
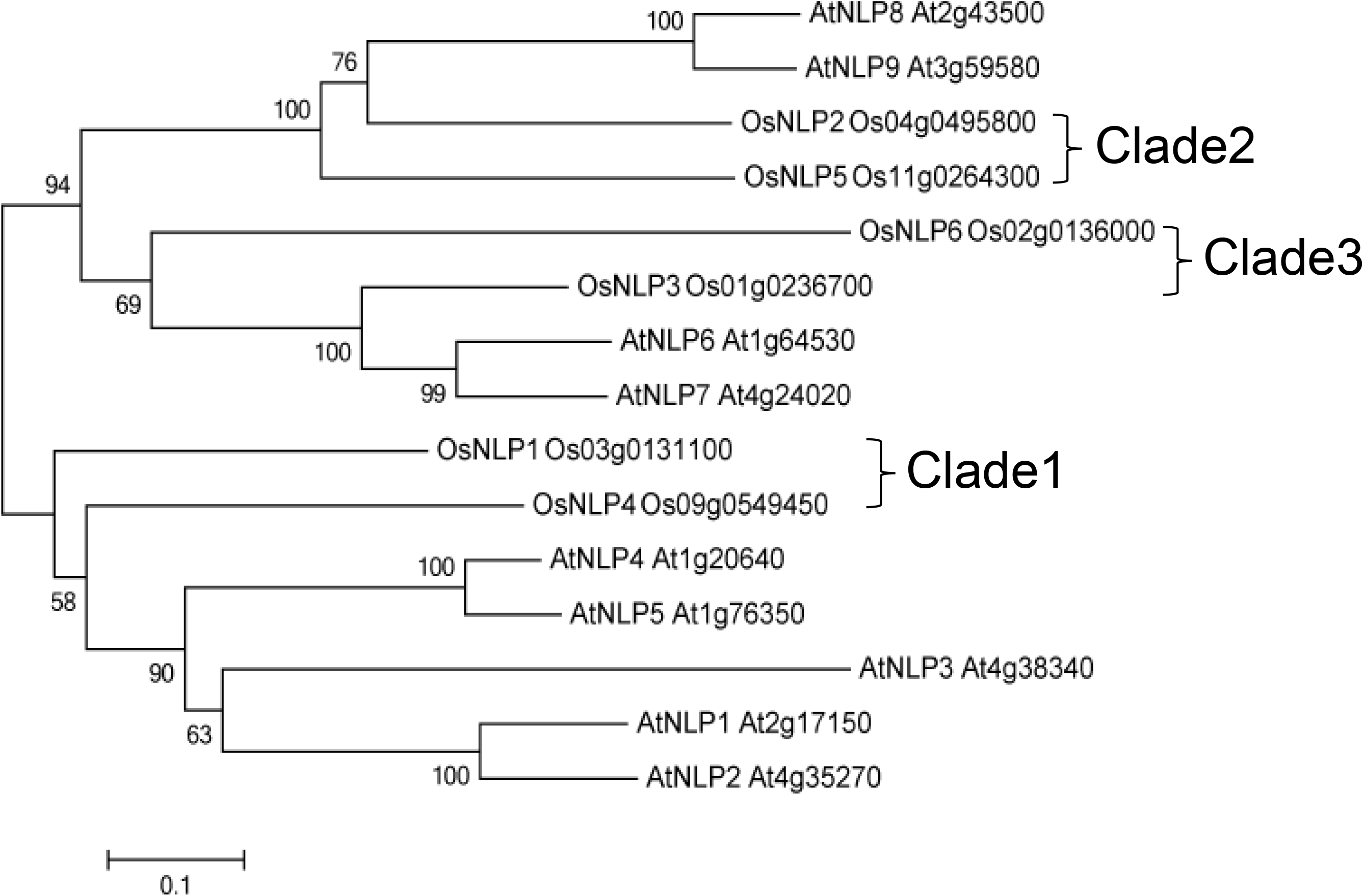
Phylogenic relationship between AtNLPs and OsNLPs. Phylogenetic tree presented evolutionary relationships between rice and *A. thaliana*. The scale bar represents the evolutionary distance. Six OsNLP family members were classified into three clades. OsNLPs sequences and AtNLPs sequences were obtained from RAP-DB (http://rapdb.dna.affrc.go.jp/) and TAIR (https://www.arabidopsis.org/) relatively. Phylogenetic tree computed by the MEGA5 software.

**Supplemental Figure S2.**
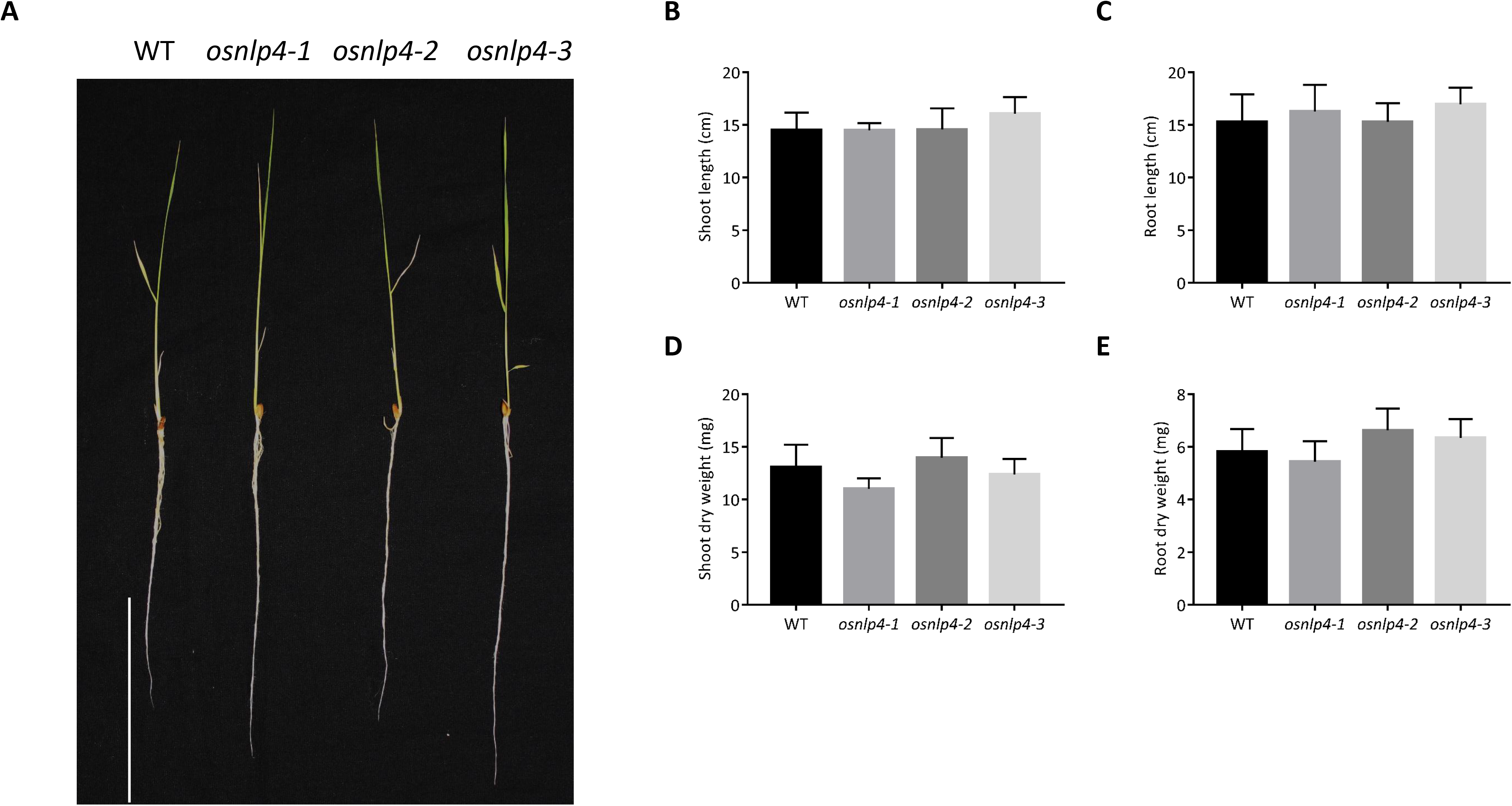
Physiological phenotype of WT and the *osnlp4* mutant lines under nitrogen free condition. **A**, Plants were grown in hydroponic culture without nitrogen for 15 days. Scale bars, 10 cm. The shoot length (**B**), root length (**C**), shoot dry weight (**D**) and root dry weight (**E**) of the *osnlp4* mutants were compared to wild type. Error bars represent the standard deviation for 6 plants. There was no statistical differences between wild type and mutants (Dunnett’s test, \**p*<0.05).

**Supplemental Figure S3.**
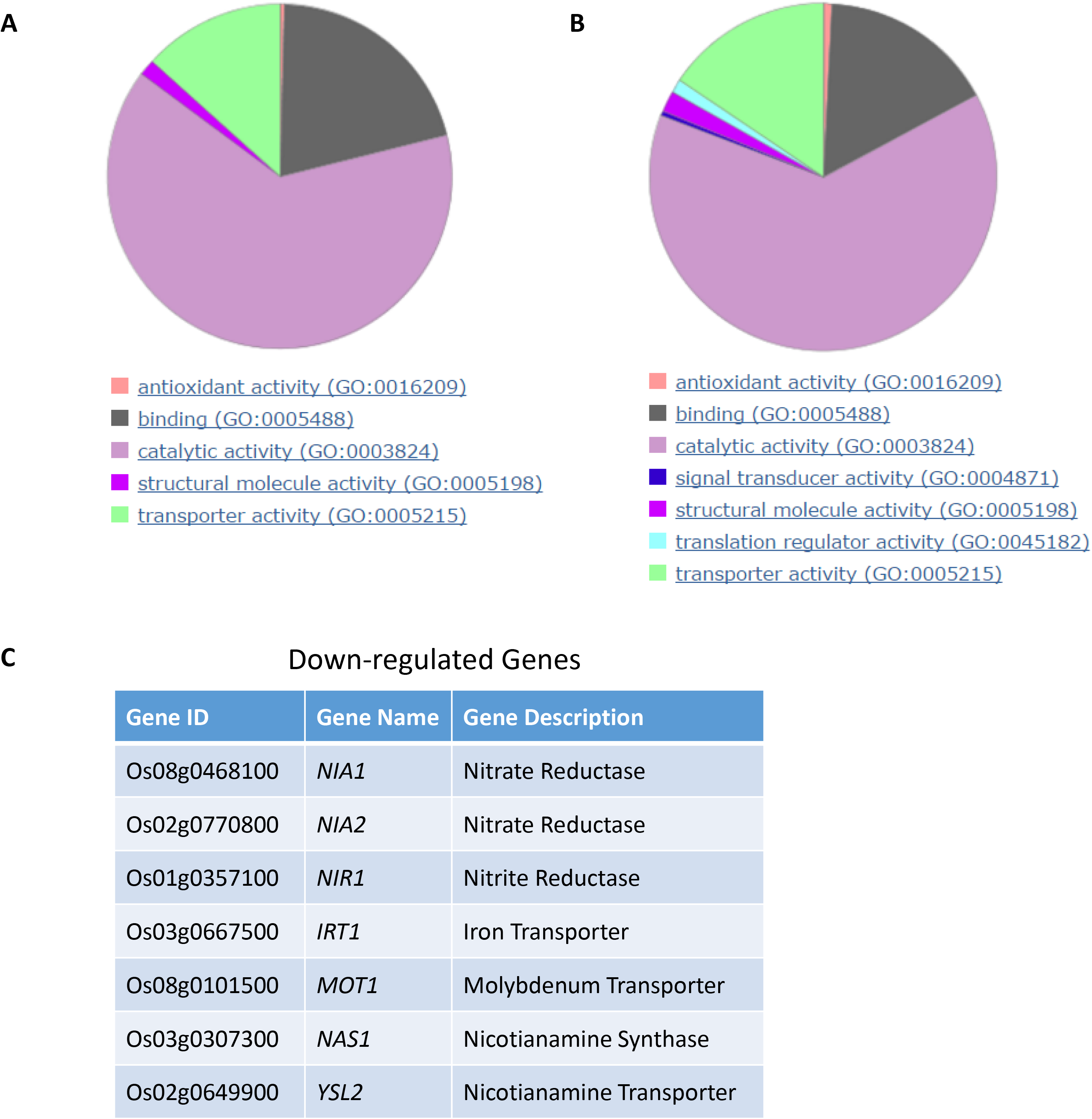
GO analysis of the *osnlp4-1* according to RNA-seq data. GO analysis showed the molecular function categories of 580 up-regulated genes (**A**) and 622 down-regulated genes (**B**) in the *osnlp4-1* under nitrate condition. **C**, A short list of down-regulated genes in the *osnlp4-1*.

**Supplementary Table 1.**
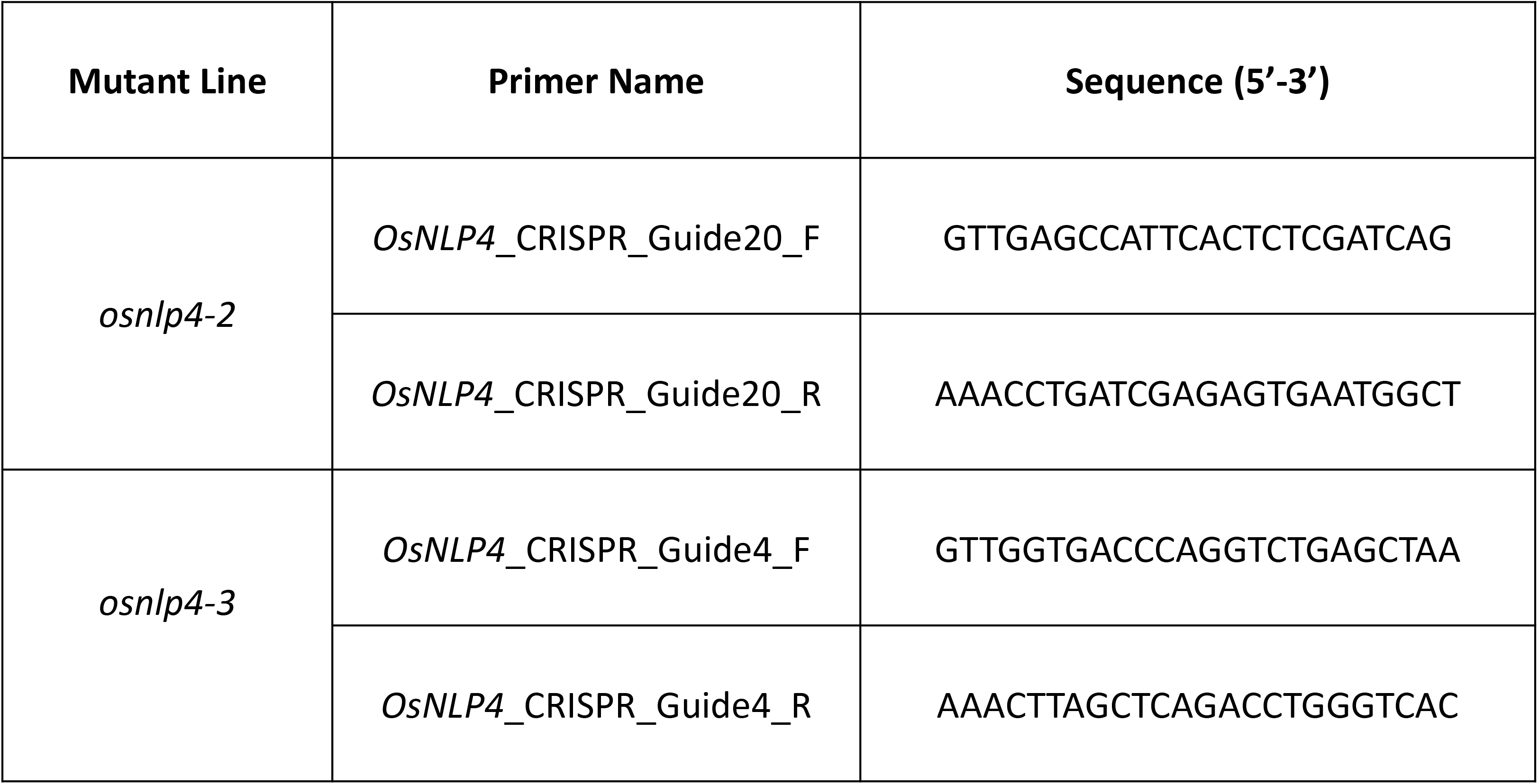
Primer pairs used for constructing *OsNLP4* CRISPR/Cas9 lines.

**Supplementary Table 2.**
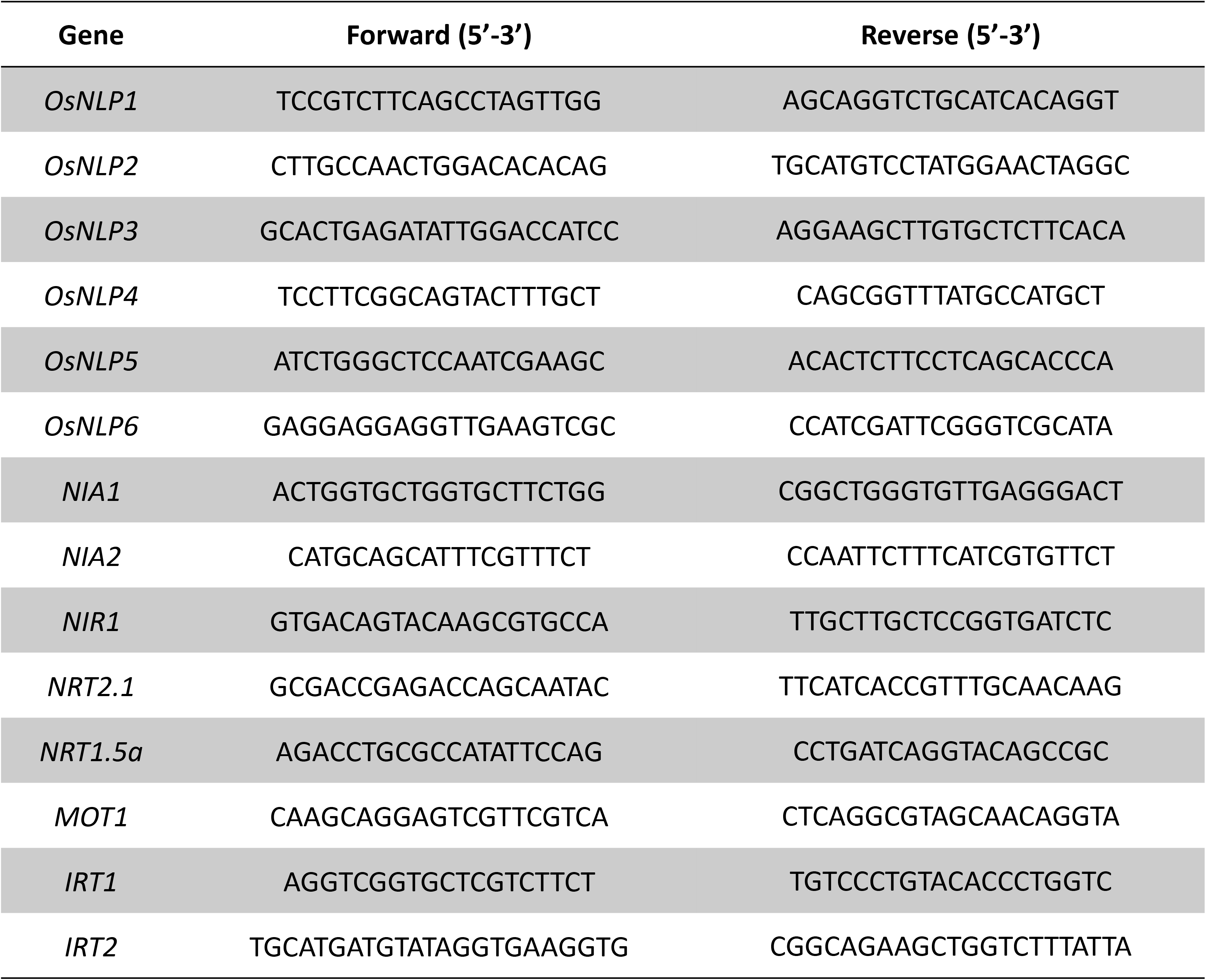
Primer pairs used for qRT-PCR.

